# Lisdexamfetamine alters BOLD-fMRI activations induced by odor cues in impulsive children

**DOI:** 10.1101/2020.04.17.046888

**Authors:** Silvia S. Hidalgo Tobón, Pilar Dies Suárez, Eduardo Barragán Pérez, Javier M. Hernández López, Julio García, Benito de Celis Alonso

## Abstract

**Introduction:** Lisdexamfetamine (LDX) is a drug used to treat ADHD/impulsive patients. Impulsivity is known to affect inhibitory, emotional and cognitive function. On the other hand, smell and odor processing are known to be affected by neurological disorders, as they are modulators of addictive and impulsive behaviors specifically. We hypothesize that, after LDX ingestion, inhibitory pathways of the brain would change, and complementary behavioral regulation mechanisms would appear to regulate decision-making and impulsivity.

**Methods:** 20 children were studied in an aleatory crossover study. Imaging of BOLD-fMRI activity, elicited by olfactory stimulation in impulsive children, was performed after either LDX or placebo ingestion.

**Results:** Findings showed that all subjects that underwent odor stimulation presented activations of similar intensities in the olfactory centers of the brain. This contrasted with inhibitory regions of the brain such as the cingulate cortex and frontal lobe regions, which demonstrated changed activity patterns and intensities. While some differences between the placebo and medicated states were found in motor areas, precuneus, cuneus, calcarine, supramarginal, cerebellum and posterior cingulate cortex, the main changes were found in frontal, temporal and parietal cortices. When comparing olfactory cues separately, pleasant food smells like chocolate seemed not to present large differences between the medicated and placebo scenarios, when compared to non-food-related smells.

**Conclusions:** We demonstrated that LDX, first, altered the inhibitory pathways of the brain, second, increased activity in large amounts of brain regions which were not activated by smell in drug-naïve patients, third, facilitated a complementary behavioral regulation mechanism, run by the cerebellum, which regulated decision-making and impulsivity in motor and frontal structures.

## Introduction

Impulsivity affects inhibitory and emotional control (1), as well as cognitive function (2). Furthermore, impulsivity is heavily related to eating disorders, specifically binge eating disorder (BED), (3, 4). Impulsivity combines a diverse set of behaviors, which can include premature decision-making, ineffective impulse control and/or inability to delay gratification (5). As there are many behaviors related to it, impulsivity should be divided into different subtypes (6). Some examples of subtypes would be non-planning impulsivity, attention impulsivity, motor impulsivity or lack of premeditation (7). Therefore, it is reasonable to expect that brain networks and brain areas will vary depending on the kind of impulsivity a subject presents with (8). Brain areas already known to show activity during impulsive behaviors are the orbitofrontal cortex (sensory integration and decision-making), cingulate cortex (inhibition) and temporal lobes (memory), as well as other areas in the limbic system, including amygdala (emotion processing and memory), insula (emotion and sensory integration), basal ganglia (memory and motivation, as well as decision-making) and thalamus (connective sensory center), (9, 10).

Several drugs are regularly used to treat ADHD/impulsivity symptomatology. Some are based on psychostimulants derived from amphetamines and methylphenidate, while other, non-psychostimulant drugs are also derived from atomoxetine and guanfacine (11). Lisdexamfetamine (LDX) has been recently added to this body of treatments. LDX is a drug used to treat ADHD/impulsive patients and patients with BED (12). It has had FDA approval since 2007 and has shown to be beneficial to patients with ADHD/impulsivity (e.g. (12)(13, 14)). It has also been proven to have similar tolerability and safety limitations to traditional treatments (15) – but work focusing on brain imaging of the effects of this drug is sparse. Fleck et al. (16) have shown that a treatment with LDX of women with BED reduced activity in the globus pallidus and ventromedial prefrontal cortex (VMPFC). Studies on menopausal women showed that the drug modulated insula and dorsolateral prefrontal cortex (DLPC) recruitment (17). In a recent study by Schulz et al. (18), children with ADHD treated with LDX were found to present weaker connectivity between amygdala and frontal cortex structures, and this reduced the emotional bias of cognitive control. Finally, studies in rodent models have shown that LDX improved ADHD-related symptoms by stimulating striatal hypertrophy (19).

The different areas of the brain which have been imaged with magnetic resonance imaging (MRI) and are related to the olfactory function have been fully described in the past (e.g. (20-22)). In general, primary olfactory cortex (prepiriform and periamygdaloid areas), amygdala, hippocampus, secondary olfactory cortex (parahippocampal gyrus, which includes perirhinal and entorhinal cortex), thalamus, insula and orbito-frontal cortex are all relevant. While primary olfactory cortex is related to detection of smells, the function of secondary cortex is related to their appreciation. Positron emission tomography (PET) studies have shown that an absence of the capacity to detect smells was related to hypoactivity of these brain regions (23). Olfaction and its brain representations are well known to be mediated through a myriad of factors, which could include genetics (24), training (25), pleasantness of a smell (26), etc. Even if a recent meta-analysis studied states that ADHD patinents present small to negligible effects on odor function crow (27), there is a large body of work which supports how specifically impulsivity, does indeed affect this function (i.e.: (28),(29),(30),(31)). From all these factors the existence of some neurological disorders are known to affects smell capacities greatly (32).

MR techniques are widely used to study neurological disorders (e.g. (30, 33, 34)); nevertheless, work linking impulsivity and odor is sparse, even when olfaction is considered to be a precursor or modulator of addictive and impulsive behaviors, (35, 36). Lesions to the orbitofrontal cortex related to impulsive aggressive disorders have shown to make patients anosmic (37). More recently, continuous theta bursts of transcranial magnetic stimulation to the right orbitofrontal cortex produced impaired odor perception (38). Olfaction is known to be a primary cue for the appearance and control of appetite and substance abuse, as odors stimulate the same brain areas as those activated by addictive substances (39). Still, the neurophysiology underlying the link between odor and impulsivity is not clear, and to our knowledge no work exists imaging impulsive children medicated with LDX while stimulated by odor cues.

The objective of this study was to image brain activity elicited by olfactory stimulation in impulsive children after LDX or placebo ingestion. Experiments were performed using functional MRI (fMRI) techniques. This protocol was completed in order to better understand the inhibitory and emotional processes of the impulsive children’s brains during olfactive stimulation.

## Methods

### a) Ethical

This study was approved by the Ethical Committee of the Hospital Infantil de México, Federico Gómez, under a project titled “Evaluation of the response of Lisdexamfetamine in children and adolescents with ADHD” (IBH code: HIM 2018-88). This study was performed in accordance with international practice, which included procedures from the Helsinki Declaration. All volunteers and caretakers in this study attended an initial information session. Once considered suitable and after agreeing to participate, they signed information and consent forms. Volunteers could abandon the protocol at any time of their choice, without giving a reason. Data from this study has been deposited in a repository and will be available under petition to the authors.

### b) Volunteers

One cohort of 20 male children between 6 and 9 years of age (7.5 ± 1.4) were recruited for this work. 20 volunteers is not a large sample, but it is comparable to samples used for other publications in the field (e.g. (30, 40, 41)). Samples this size have been shown to have enough statistical power to show significant results (42). A Tanner test was performed to make sure none had reached adolescence. Volunteers were recruited from normal walk-in patients to the neurological clinic of the hospital. All children were right-handed. Volunteers did not have a history of neurological disorders (e.g. epilepsy, autism, cognitive dysfunction, etc.), and they could participate in MRI procedures. Body mass indices of the volunteers averaged 28.0 ± 2.7, which placed them mainly in the overweight and obese range (n=4 were normo-weight). Volunteers were screened for eating disorders. None presented any.

As the fMRI study performed on volunteers in this study included olfactory stimuli, normal smell detection and interpretation had to be assessed for all volunteers. First, normosomiay of volunteers was assessed by an otorhinolaryngologist. Then preliminary bench studies were performed with volunteers lying in the scanner to assess that they were able to detect and differentiate the three smells provided during scans in the amounts provided by the protocol. To conclude preliminary preparations, a series of questions on the hedonic value of the smells was performed on volunteers, to make sure that no “strange” response to a specific smell could be elicited.

The main objective of this project was to assess effects on physiology of smell driven by impulsivity. Nevertheless, all the volunteers that participated in this study had ADHD of the hyperactive-impulsive subtype. Because of this, even if not relevant to the findings and results of this study, a brief description on how their diagnosis was made is given. ADHD in patients was diagnosed by specialists in the field (neurologists and psychiatrists) after clinical interviews with and without the presence of parents and people in charge of infants. These interviews were completed with evaluation of volunteers through ADHD and Aula tests. In order to “quantify” the severity of ADHD of volunteers for this study, data according to the DSM-5 criteria was used. Time from diagnosis for volunteers in this study was short (1.3 ± 0.3 years), and all were drug-naïve. Furthermore, through a detailed study of clinical records, volunteers with a known hypersensitivity to LDX components, or comorbidities known to be accentuated by the drug, were excluded from the study. Impulsivity was assessed with a Go/No-Go task, which is a gold-standard for assessing this trait (e.g. (43, 44)). The results of the test were not quantitative, and only indicated if the patient was or was not impulsive. All volunteers showed the impulsivity trait for this study. It is worth mentioning that this test is unable to subdivide impulsivity into different sub-traits. In some cases, a complementary MOXO-dcpT fMRI scan was performed (45). This test is not an integral part of ADHD diagnostic procedures, but it can be used to assess impulsivity.

### c) Paradigm

Volunteers for this study visited the research center on three occasions. During their first visit, anthropometric measurements, ADHD and impulsivity as well as clinical evaluations were performed. If volunteers fulfilled all inclusion criteria, they then visited the MR unit on a second and third day. The paradigm and imaging protocol followed was the same on days two and three. The only difference was that on one day patients ingested LDX, and on the other day they ingested a placebo. The order in which placebo or LDX was given to the volunteers was random, making sure that 10 patients got placebo the first day and the other 10 LDX. This was therefore an aleatory crossover study. On each visit, all imaging was performed between 7 and 11 am in the morning, and patients were asked to have had a light breakfast (they were not fasted). First, placebo or drug were delivered to the patient with him fully dressed and out of the scanner. The medication used in this study was LDX (Vyvanse®), which was administered orally in a one-time dose of 15 mg per volunteer. After 2 hours’ wait, during which volunteers remained in the clinic in a playroom, they performed a Conners’ Global Index test (CGI test) (46) under the supervision of a trained neurologist. Volunteers were then asked to prepare and lie inside the MR scanner. Once there, volunteers underwent MR sequence preparations, an fMRI study (blood oxygen level dependent imaging, BOLD) and anatomical imaging. The whole imaging process took a total of approximately 20 minutes. Once the three repetitions of this protocol had been completed, participation in the study was considered to have finished.

The BOLD-fMRI experiment used three odor cues which were randomly presented (chocolate, clove and lavender). Chocolate represented a high calorie content with a large hedonistic value, clove represented healthy foods with a neutral hedonistic value and lavender a food-unrelated smell with a more positive hedonistic value, (47, 48). Odors were presented to the volunteers by a technician situated behind the scanner, who was cued by a video presentation shown on a screen visible exclusively by him/her. Volunteers were instructed not to move during the fMRI scan, especially their noses and mouths, to reduce motion artifacts in the results. A detailed explanation of this paradigm is provided in recent research from this group (30).

### d) Hardware

Imaging was performed on a 1.5 T Philips-Intera Achieva scanner with a NOVA gradient system set. A birdcage head coil with SENSE technology and 8 channels was used for fast imaging. BOLD imaging consisted of the acquisition of 278 brain volumes with TR=3000 ms. A T_2_*-weighted gradient-echo sequence was used with TE=30 ms, flip angle=80°. 35 consecutive axial slices were acquired (without gaps) and in ascending order, covering the whole brain from frontal lobe to cerebellum. An image resolution of 3.05 mm in plane and 4.5 mm thick slices were obtained with 80×80×35 matrices. Anatomical images matched the position and volume studied by the BOLD imaging and were obtained with a fast T_1_-weighted gradient-echo sequence (TR=307.81 ms, TE=2.48 ms, flip angle=80°). Resolution for these images was 0.38×0.38×4.5 mm with a 640×640×35 matrix. Four whole-brain acquisitions were obtained and averaged (NE=4).

### e) Software and Image Analysis

Image analysis of this data was identical to the methodology described in previous work from this group, (30, 40, 41). Briefly, BOLD-fMRI data was preprocessed with the CONN toolbox (49). This toolbox was run on a Matlab platform (Mathworks, Natick, MA, USA) version 2017a. On top of this, SPM12 (www.fil.ion.ucl.ac.uk/spm/) was available in the background, as its subroutines were used by the CONN toolbox. Pre-processing used CONN’s standard analysis pipeline and included the following steps:

1. Realignment and unwrapping.
2. Slice time correction to the middle slice.
3. Segmentation of CSF, white and gray matter in anatomical images.
4. Normalization of anatomical brain image to MNI brain template.
5. Normalization of fMRI data to MNI brain template.
6. Functional outlier detection with the Automatic Registration Toolbox (ART) tool (https://www.nitrc.org/projects/art/).
7. Smoothing with a Gaussian kernel 3 times the size of the voxels (FWHM).

After pre-processing, data was fed to SPM12, and a first-level fMRI general linear model (GLM) was built using all the MR data (scans after LDX or PLA intake). In it, four different contrasts were considered: one per smell (clove, chocolate and lavender) and one considering all the stimuli together (all smells). Second-level contrasts were built in the GLM to assess the differences for the same four contrasts, but comparing the placebo (PLA) and LDX scenarios.

The significance of images obtained here was established as clusters of at least 10 voxels with a p<0.01 after family-wise error (FWE) correction calculated using random field theory. The threshold selection is always a controversial choice and much discussion exists on the ideal value to be used. The values selected here are standard values, common to other publications in the field (e.g. (30, 41)). Images were presented using the Viewer option of the DPARSFA software (50), in which fMRI-calculated t-maps were overlaid on anatomical MNI brain templates, using the statistical threshold mentioned. Montages in the axial direction were then created.

Finally, numbers of activated voxels in each cluster as well as the region in which they were found was extracted. Regions are presented with both the Automated Anatomical Label (AAL) atlas and Brodmann area (BA) information. When lateralization was calculated for this work, a proportion (%) of voxels activated in each hemisphere with respect to the total was calculated.

## Results

Psychological evaluation showed a reduction in CGI scores from 6.5 ± 1.1 (mean ± SD) for the PLA state to 2.4 ± 0.9 in the LDX state. This difference reached statistical significance after performing a paired t-test (p=0.008).

Data (in the form of t-maps) from the fMRI-BOLD experiment in which all smells delivered to patients were considered together are presented in **Figure 1**. Here information for the PLA and LDX states are presented together with the LDX-PLA contrast. This is done to assess the hypothesis presented in introduction, which states that after LDX ingestion inhibitory pathways and behavioral regulation of the brain change after olfaction cues to regulate decision making in impulsive children. Activations were larger in volume for the LDX images when compared to the PLA situation: there was a ratio of almost 5 to 1, as can be observed in Figure 1 (53926 voxels vs. 11048). Lateralization was also found to increase with LDX. In the PLA state, 55.8% of activated structures were in the left hemisphere for patients, increasing to 60.3% after medication.

**Figure 1.**
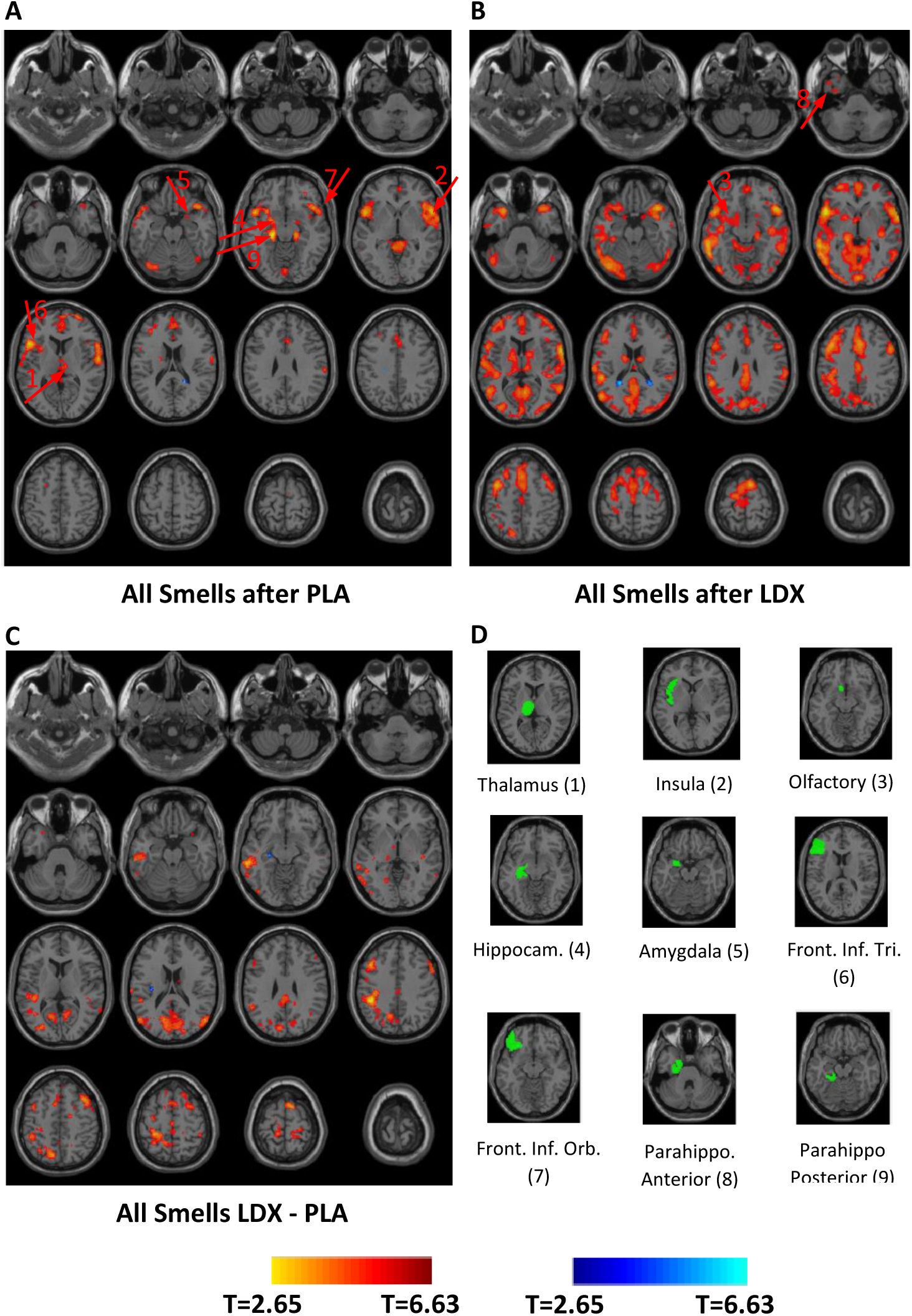
BOLD-fMRI results for all odor cues used in this study together. This figure presents BOLD-fMRI activations for the three stimuli used in this study together (chocolate, clove and lavender). Results are presented on 16 axial slices (2 cm thick) that cover the whole brain. **A** presents activations for the PLA experiment. **B** presents the results after LDX ingestion. **C** presents the results for the LDX – PLA contrast. Significance of the results is presented with a pseudocolored scale in which hot colors (red, orange) represent large significance or correlation with the stimuli. Cold colors (blue and white) represent anticorrelation with stimulus in **A** & **B** or stronger activation for the PLA state in **C**. A t-score scale is added under the images to quantify significance of results. Panel **D** presents nine regions related to perception of smell. They are numbered and correspond to the red numbered arrows on **A** and **B** pointing at these specific activations. This image shows that no significant differences were found between PLA and LDX states in olfactory perception centers of the brain. This can be seen by the absence of activations in these regions in the LDX-PLA contrast (**C**). Differences were found in contrast in: frontotemporal and parietal areas of the brain, together with regions like cingulate cortex, motor areas, cerebellum and posterior cingulate cortex.

A whole list of the areas activated in the PLA state can be found in **Table 1**. There were nine areas activated in the frontal cortex, including the inferior opercular area. This contrasted with just one in the temporal and occipital cortex respectively. Activation in areas related to olfaction was also found (hippocampus (bilateral), parahippocampal, insula, thalamus, olfactory cortex, amygdala and frontal inferior orbital). Structures related to motion were also activated (precentral and superior motor areas (bilaterally)), as well as a series of relevant structures such as cuneus, caudate, anterior cingulate cortex (ACC), medial cingulate, calcarine, cerebellum crus I and the rolandic operculum. The first three of these latter areas are known to be affected by ADHD.

**Table 1:**
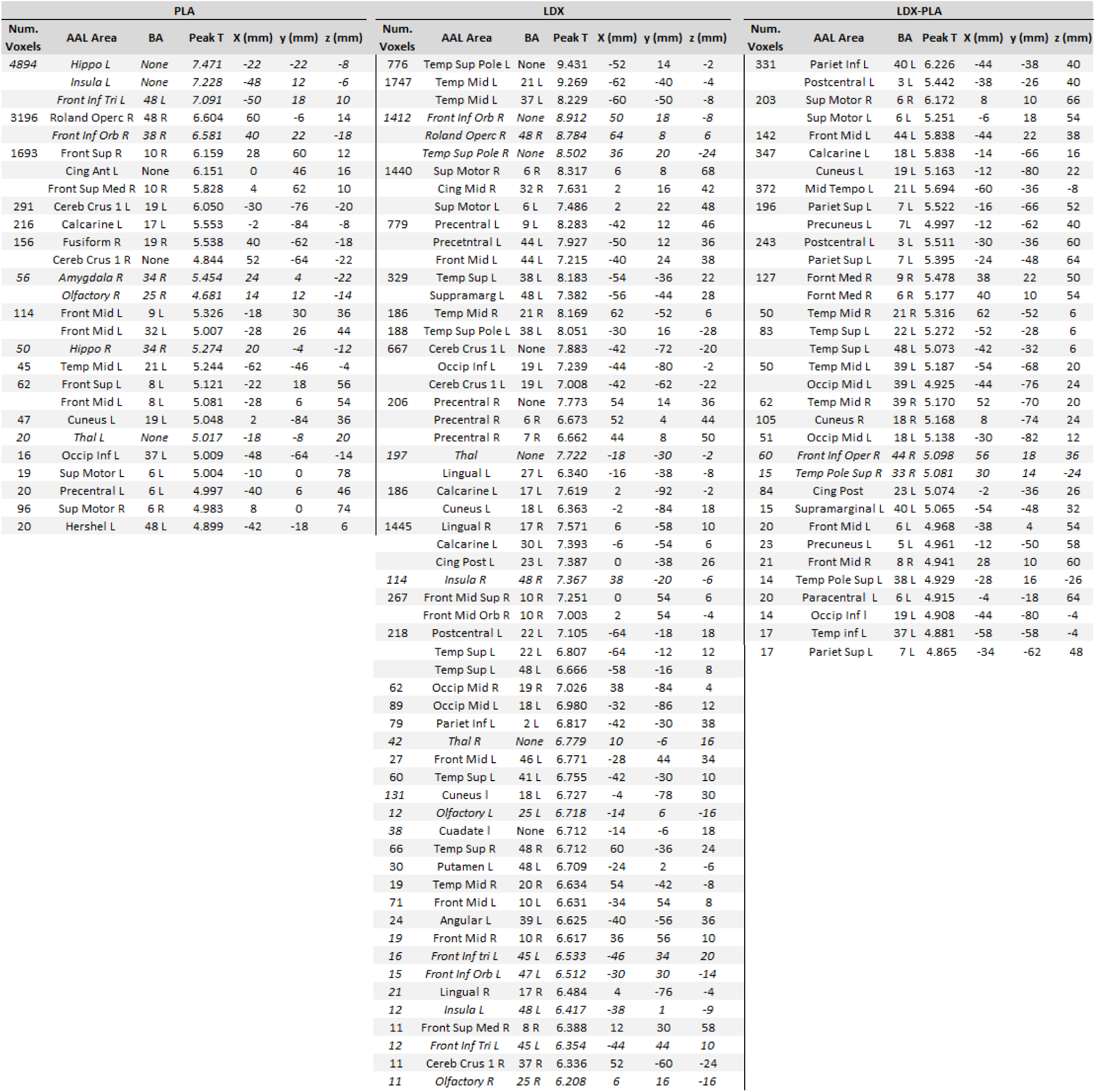
Quantification of BOLD-fMRI results for all odor cues together. Quantified data presented for all olfactory stimuli together for the following contrasts: PLA, LDX and LDX– PLA. For each stimulus, first the number of activated voxels in each significant cluster is given. When a number of voxels is not followed in the following row by other number, it means that the same cluster was large enough to have different maximum points. After this are given the regions that these clusters were in. This is done showing both the corresponding AAL region and the BA in which they appear. In the last four columns, the statistical significance of the maximum voxel is presented, together with its MNI coordinates in mm. Regions in italics in this table are regions directly related to olfactive perception.

For the LDX case, twelve areas were activated in the fontal cortex, including the inferior orbital which was activated bilaterally. In contrast with the PLA scans, ten different activations appeared in the temporal lobe (the majority, 70%, in the superior lobe) and three in the occipital lobe. Activation of areas related to olfaction reproduced the ones found before, with two major changes. First, the amygdala was not found to be activated in this LDX case and, even though all structures related to olfaction now appear with bilateral activations, in contrast with the previous scenario, activation of the hippocampus is much less significant than before. Structures related to motion were also activated (precentral and superior motor areas, both bilaterally). Second, a series of relevant structures such as cuneus (three times more), caudate, calcarine, cerebellum crus I (2 times more and bilateral) and the rolandic operculum showed activations too. Cingulate cortex activations changed from the previous case. The whole cingulate cortex presented activations, in contrast to PLA with activations in the anterior and minimal ones in the medial. New regions were activated in this case, namely lingual and angular gyrus.

All olfactory structures seemed to present similar strength and volume of activation in both states, as no significant differences between them were found. This can be seen by the lack of activation of olfactive structures for this contrast in Table 1 (only marginal activation in frontal inferior cortex). There was a trend in the LDX states to present larger activations in the precentral and supplementary areas. These activations were nevertheless found before in both states. The main difference for this contrast were the larger activations in the frontal (BA 6, 8, 9 and 44), temporal (BA 21, 39 and 48), parietal (BA 7 and 40) and occipital lobes (BA 18 and 39) for the LDX state. Regions with larger activations in the PLA state included the calcarine, cuneus, angular, precuneus, and supramarginal areas. Finally, the cingulate cortex presented significant differences in activation, but only in the posterior area.

Data (in the form of z-maps) are presented in **Figure 2** for the fMRI-BOLD experiment, in which each smell (chocolate, clove or lavender) is considered separately, and are presented exclusively as a LDX–PLA contrast. The aim of this part of the study was to find differences in odor responses due to the distinct hedonic value of the smells. We wanted to assess if impulsivity (with its alleged different inhibitory mechanisms), treated differently smells as chocolate which can drive excitement, hunger and similar responses, compared to lavender which is not even a food related smell. In the case of chocolate there were a larger number of activations in the PLA situation than for the LDX. This difference was marginal (235 voxels vs. 197). For clove, larger activations were found LDX than PLA (8419 vs. 122 voxels). For lavender a similar situation to clove was observed, with 7682 voxels activated in the LDX state vs. 220 in PLA. Lateralization was also studied for each smell in the LDX-PLA contrast. Clove had 68.7% of structures activated in the left hemisphere of the brain while lavender had 75%. The number of significant activations in the chocolate contrast was so small that differences in lateralization for this contrast were not considered to reach significance. Nevertheless, 67% of activated structures were in the right hemisphere, in contrast with the other two smells.

**Figure 2.**
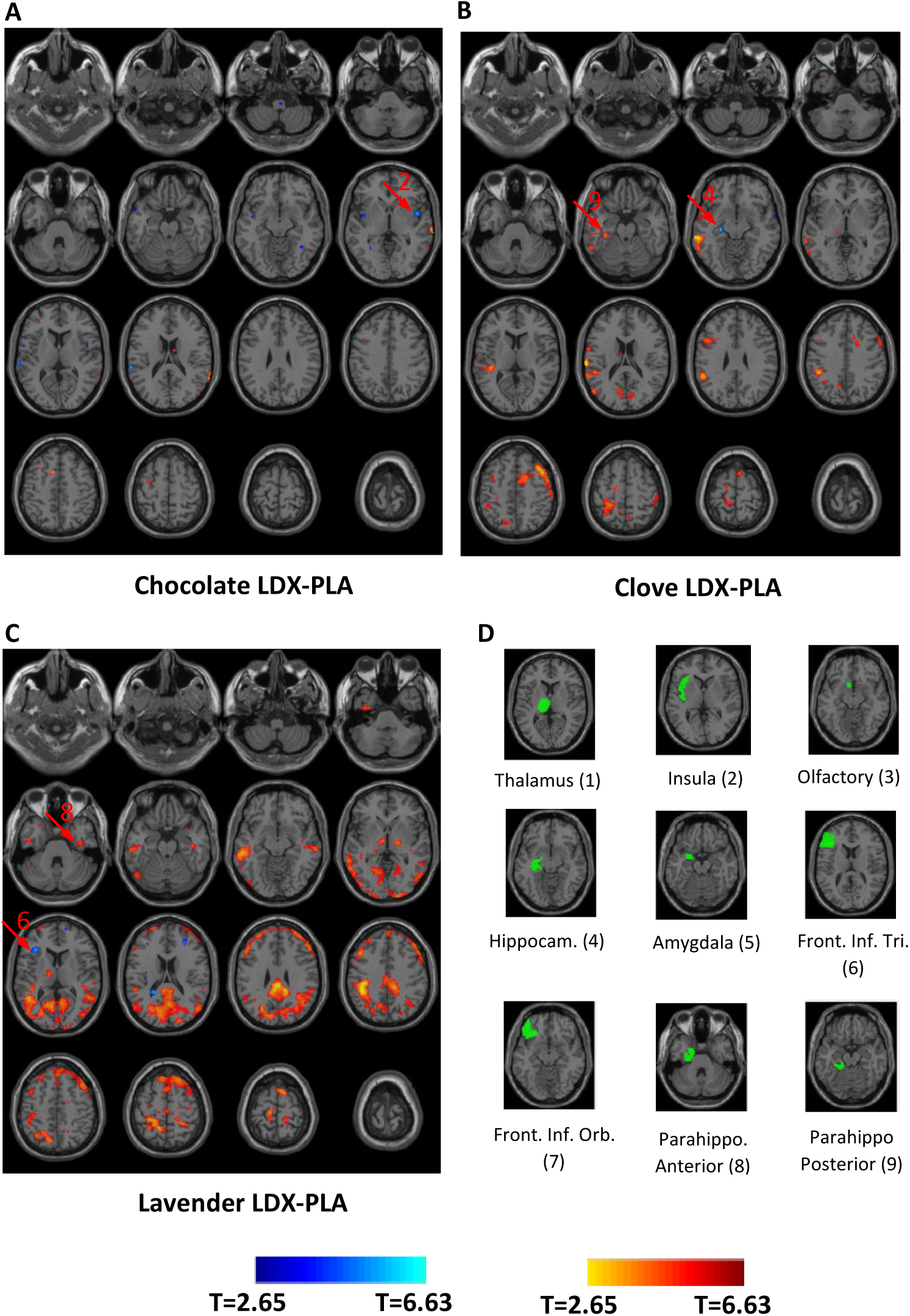
BOLD-fMRI results for each odor cue for the contrast LDX–PLA. This figure presents BOLD-fMRI activations for each one of the three stimuli used in this study: chocolate **A**, clove **B** and lavender **C** for the LDX – PLA contrast. Results are presented in the same format as Figure 1 (see legend from Figure 1). This figure shows that pleasant food smells like chocolate seemed not to present large differences between the medicated and placebo scenarios when compared to other non-food-related smells.

A whole list of the areas activated for each smell in the LDX-PLA contrast can be found in **Table 2**. Chocolate flavor, which produced the smallest number of activations in the LDX–PLA contrast presented positive activations in middle frontal lobe (bilateral BA 8, 11 and 44), the superior temporal lobe (BA 21, 22 and 48) and the left superior motor area. In contrast, negative activations (PLA state “stronger” than LDX) were found in insula (bilateral), fusiform (bilateral), the superior temporal cortex and the frontal inferior orbital cortex. Clove presented much larger activations than chocolate. A main body of activations was found in the middle temporal cortex, with ten clusters of activity, followed by five in the superior temporal cortex. These results were completed by activations in the frontal cortex (one in each area: superior, middle and inferior). Almost no activations were found in the parietal and occipital lobes. Some positive activations in the LDX situation, when compared to PLA, were found in structures related to olfaction such as the hippocampus, insula, thalamus, frontal inferior orbital and parahippocampal cortex. There were none in the olfactory cortex or the amygdala region. Large differential activations were found in cuneus (bilateral) and precuneus, calcarine (bilateral), supramarginal (bilateral), fusiform and in the rolandic operculum. Precentral, postcentral (three clusters) and supplementary motor areas were all activated bilaterally for this smell. Finally, activation in cerebellum was found for this smell in crus I and in cerebellum 6. Two negative activations appeared, in hippocampus and the superior temporal lobe.

**Table 2:**
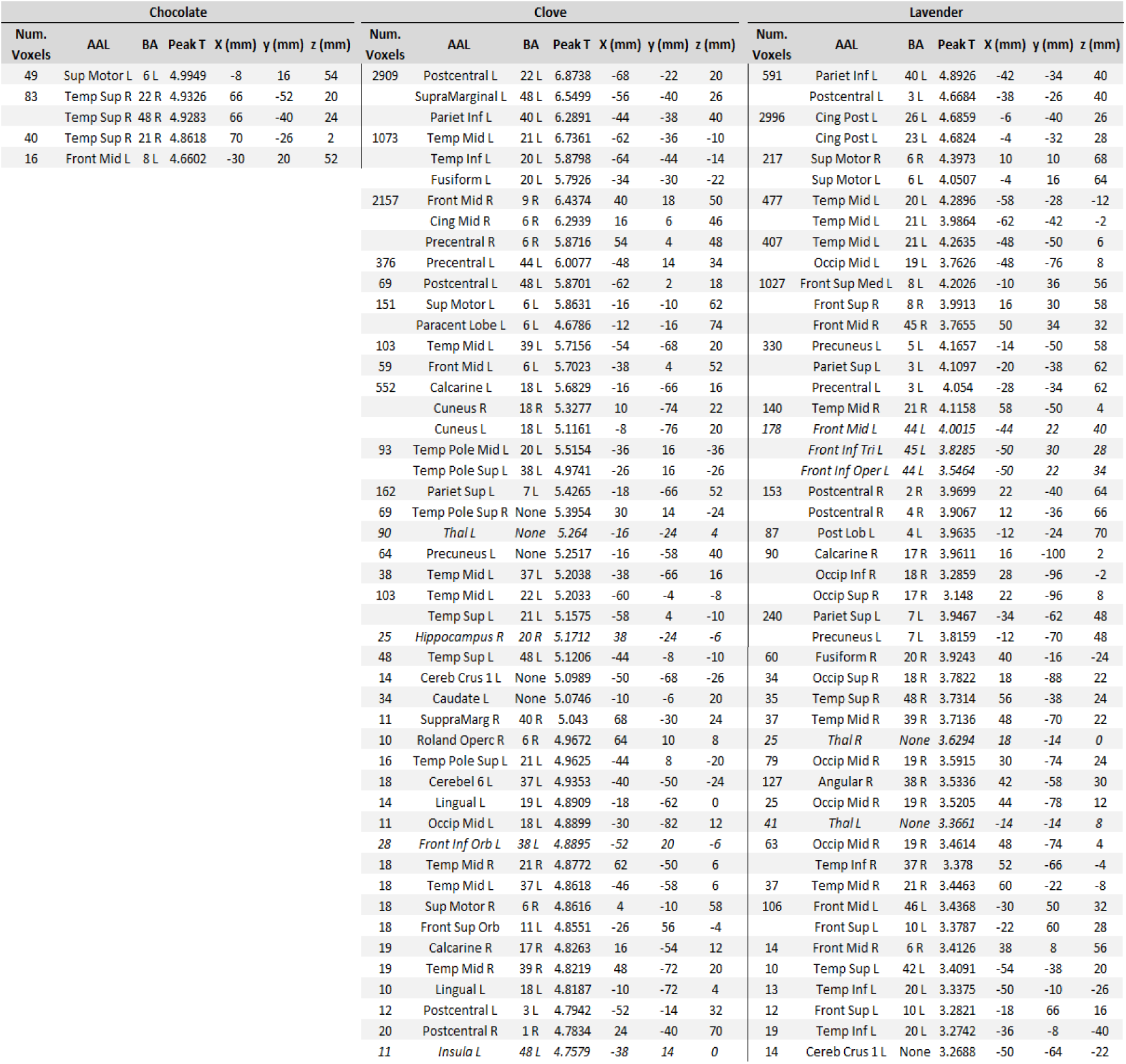
Quantification of BOLD-fMRI results for each odor cue for the LDX–PLA contrast. Quantified data is shown from regions activated for the three contrasts presented in Figure 2. Data is presented for all olfactory stimuli separately, for the following contrasts: chocolate LDX–PLA, clove LDX–PLA and lavender LDX–PLA. Format of table is identical to that of Table 1 (see Table’s 1 legend).

For the lavender smell, and in contrast to clove and chocolate, a large body of activations were found in the occipital cortex (two clusters in the superior, four in the middle and one in the inferior). As in the clove case, there were activations in the frontal areas, which were more numerous in the middle and superior lobe. The temporal lobe was still the one most activated by smell and had activations in superior (3) middle (5) and inferior (1) lobes. These were approximately 50% of those seen for the clove stimulus. Here, a small number of structures related to olfaction were activated, with thalamus and the inferior frontal lobe being the only ones presenting differential activations. Some relevant areas activated here were the precentral and postcentral (bilateral), fusiform, angular and precuneus areas. In contrast with the clove smell, no cuneus, calcarine or caudate activations were found. Some activation in the cerebellum crus I was found and activation in cingulum migrated from medial to posterior. In contrast, negative BOLD results appeared in the frontal inferior orbital, insula and medial frontal lobe.

The contrasts for each smell with LDX or PLA are presented as **Figure 3**. It is of interest to see that chocolate was the contrast with the largest number of activations, both in the PLA-state (with 30525 activated voxels) and after LDX (with 37511 voxels, representing an increase of 22.9%). In contrast, clove presented 7427 activated voxels in the PLA-state, vs. 46511 after LDX-medication, an increase of 626%. Lavender in the PLA state showed 10959 activated voxels, and in the LDX state 69496, representing an increase of 634%.

**Figure 3:**
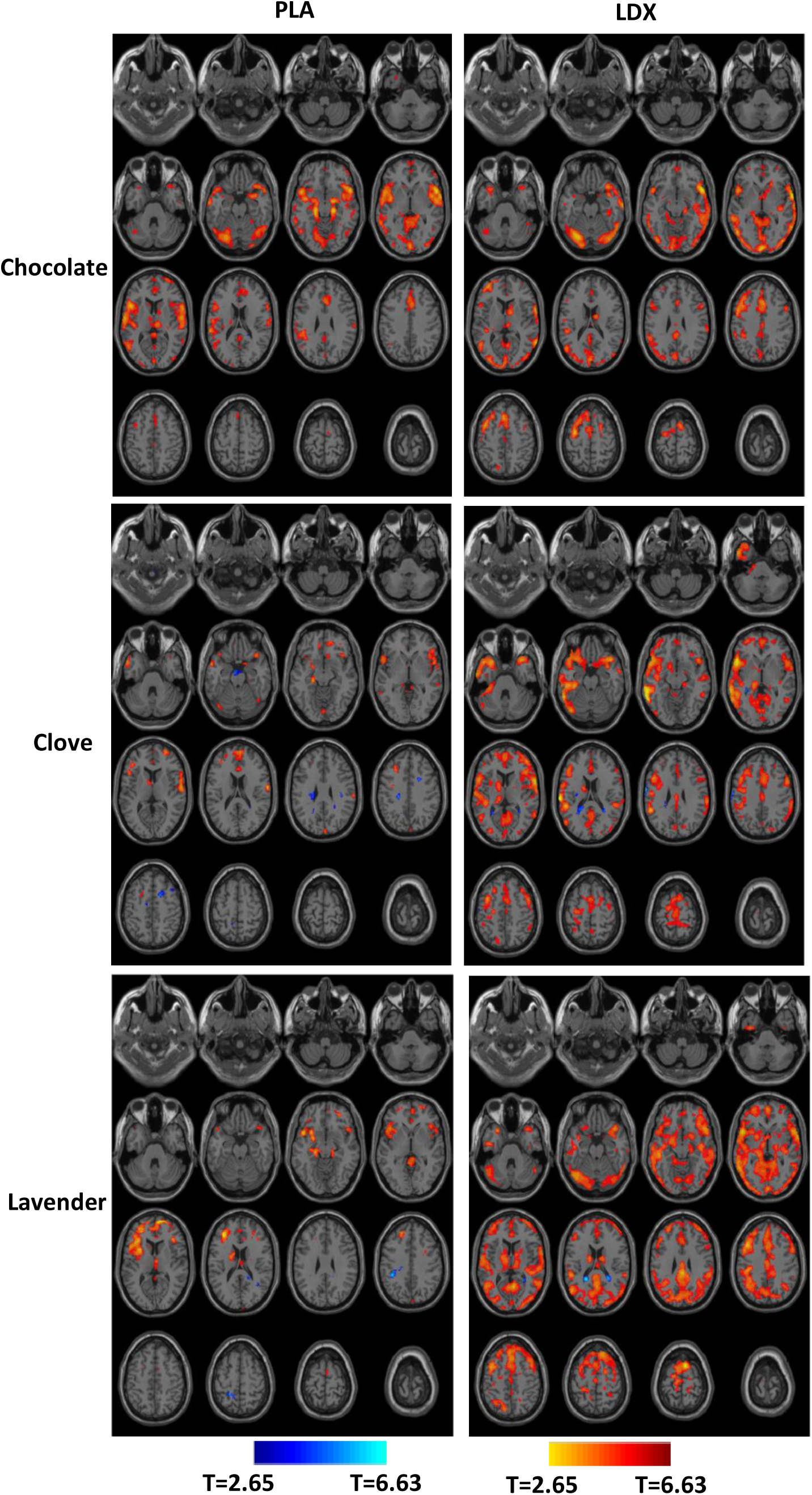
BOLD-fMRI results for each odor cue for PLA or LDX. This figure presents BOLD-fMRI activations for each one of the three stimuli used in this study: chocolate, clove and lavender. Results are presented in the same format as Figure 1 (see legend from Figure 1) with the exception of the ROI related to odor perception of panels **D**. It can be observed that LDX increased brain activity in general when a flavor did not entice it on its own in the PLA group,

## Discussion & Conclusions

The main findings of this paper were as follows. First, impulsive patients after receiving a dose of LDX showed decreases in their CGI scores. Second, intake of this drug increased BOLD activation and its lateralization (towards the left) during olfactory fMRI stimulation. Third, when subjects received odor cues, activations in the standard olfactory centers were found as expected. Fourth, differences between the PLA and LDX states for all olfactory cues considered together were found in motor areas, precuneus, cuneus, calcarine, supramarginal and posterior cingulate cortex (PCC). Also, large differences in activations in frontal, temporal and parietal cortices were found. Finally, when comparing olfactory cues separately, pleasant food smells like chocolate seemed not to present large differences between the LDX and PLA stimulations, when compared to non-food-related smells like lavender or healthy food-related smells like clove.

### Limitations of the study

This study is limited by several factors. First, a sample of 20 volunteers is valid for most imaging work but not optimal, as a larger population could be used to obtain more significant results. Children undergo substantial changes between the ages of 6 and 18 years, while passing from infancy to adolescence. Because of this we used Tanner’s tests to make sure adolescent development had not started in our volunteers (ages 6 to 9). Even though this course of action has given us positive results in the past, follicle stimulating hormone measurements could ideally have been performed to make sure patients were not adolescent. The fact that no olfactometer was available to perform the study is also a limitation. Stimulus delivery might have been imperfect due to this fact, delivering different intensities of smell and during different times. To compensate for this, larger numbers of odor stimulation cues were delivered to volunteers, and exhaustive training as well as preliminary tasks were performed on them to reduce possible sources of noise. Ideally and to complete this study, connectivity measures between different brain regions would have been performed. Unfortunately, the software used here did not allow us to study connectivity between regions of the cerebellum.

### Lateralization

Understanding the lateralization of brain activity is of interest as it carries inherent information on which brain areas are active and how they communicate to regulate processes. It has been reported using fMRI that right-hemisphere structures are more dominant for right-handed humans (51) and animals (52). Lesions to this hemisphere decrease the ability to detect and distinguish smells more than to the left hemisphere (53). Finally, it has been demonstrated with fMRI that pleasant smells activate more left-hemisphere structures than unpleasant ones, which activate right-brain structures (54). In view of the findings of this study, it could be argued that our results mirror this last study, as odor cues (clove, lavender and chocolate) were all positive on a hedonistic scale and presented 69, 75 and 67% of laterality in favor of the left structure associated with pleasant smells. In ADHD, there is great body of work considering lateralization. There is nevertheless a tendency to consider that there is a right hemisphere dysfunction modulated by comorbidities associated with the disorder (55).

In our results, laterality was increased by LDX medication, not reduced. This could be seen to contradict previous findings (51, 54), but needs to be considered with care. First, brain regions related to olfaction detection presented some degree of laterality before medication but not after it (see Table 1). So, detection and basic odor processing was affected positively or as expected by LDX intake. Regions of the brain which presented greater lateralization after LDX intake were parietal and particularly temporal structures. There is a clear increase in regulation of impulsive behaviors through the self-perception and emotional association capacities of these brain areas. (16, 19) We hypothesize below that this effect could be mediated through cerebellar function. In contrast to temporal and parietal structures, frontal right regions, especially Brodmann areas 8 and 9, which are associated with the DLPC (17) and its inhibitory function, started presenting stronger activity after LDX intake. This is an effect which could be expected.

### Different strategies for odor processing were found after LDX ingestion

One of the main conclusions that can be derived from this work is that there was a clear switch of strategies and brain areas recruited for odor detection and integration after LDX consumption. The primary and secondary olfactory regions seemed to be unaffected by the drug and elicited similar responses in strength and duration. Differences in activation were more notable in other brain structures. Recent work has hypothesized that the ACC plays a crucial role in olfaction attention through links to the anterior olfactory nucleus (56). In this study, activations were found in the olfactory nucleus in both scenarios (LDX and PLA). Smaller activations were also found in the ACC in both scenarios after medication. This would support the theory, and indicates that LDX altered this modulation mechanism. If the cingulate cortex is considered, there was a shift in activations from the anterior/medial region to the posterior areas during olfaction. This is interesting as, before LDX administration, hyperactivations in anterior cortex related to inhibition, attention and decision-making functions were found. In fact, larger activations have been reported for impulsive (vs. control) groups, during fMRI experiments in adolescents who performed Go/No-Go tasks (57). These findings corroborated these results but contrasted with those of Bush et al. which found the opposite effect, with smaller activations for impulsive patients in the ACC when compared to controls during counting tasks (58). Even though some of the results presented here corroborate some previous work, there is a clear controversy in the field with respect to the role of the ACC.

The PCC is known to have a relevant cognitive role in behavior, even if there is no consensus on how this happens (59). It is a part of the default network and plays a significant role in human awareness through its links to the cuneus and precuneus. In the impulsive brain it alters the deactivation of the DMN, reducing the individual’s capability of focusing on a task (60). This effect can be mediated and brought back to normal after medication. Alterations of connectivity between PCC and cuneus have been used as an indicator of ADHD (61), and suppression of its activity reduced mind-wandering and increased attention on a given task (62). In the odor scenario it was found that greater activity was found after medication in cuneus and parietal lobes, close to PCC. This was a clear indicator that LDX was shifting away odor processing from the “abnormal” function of the ACC towards a more focused one in PCC in which attention was not lost and episodic memory retrieval played a key role.

The cerebellum has stopped being considered just as a repository region of the brain with instructions for complex movements. It is now known to play a role in control and regulation of impulsivity and other cognitive processes (63), including decision-making (64). Furthermore, it is known to play a role in cognition and emotional processing (65). Crus I and II have been found to be involved in odor processing, including detection (66) and discrimination of intensities as well as kinds of odor (67). A general trend of these experiments was finding activations in cerebellum and motor areas for odor stimuli. This happened in both scenarios (PLA and LDX), but the effect was more prominent after LDX ingestion, when larger activations in premotor, supplementary and cerebellum areas were also found. The new functions for cerebellum introduced here might have found an echo in our findings. First, the cerebellar activations discovered here mainly appeared in crus I bilaterally after LDX medication. Therefore, they were showing stronger smell detection and processing in a “normal manner” after LDX ingestion. Second, previous research has hypothesized that the cerebellum plays a pivotal role in decision-making in impulsive children (30). This happened through a possible connection with the precuneus, which modulated decision-making. Precuneus and cuneus were indeed regions which largely augmented their activation after LDX ingestion. After LDX ingestion, the frontal areas activated transitioned from being almost exclusively in DLPC to include DLPC and regions in Broca’s Area. DLPC is a region which could be involved in inhibition and abstract planning, and after medication plenty of activation still appeared in this area. But Broca’s area activation appeared only in Brodmann area 44 after LDX in the LDX–PLA contrast. This region, which is commonly associated with speech, has also been recently related to inhibition by suppression of impulsive behaviors (68). All the factors presented here pointed toward a complementary behavioral regulation mechanism run by the cerebellum, which regulated decision-making and impulsivity in motor and frontal structures: a complementary mechanism which was facilitated by LDX intake.

### Different odors elicit different responses

With respect to findings from Figure 2 and Figure 3, in previous work of this research group de Celis (30). Activations induced by chocolate were assessed comparing a control to an impulsive group. This for children older than those presented here (ages between 10-15 years). It was found that there were significant changes in connectivity between groups but changes in BOLD activations were not that large. They were only apparent in insula, cerebellum and cingulate cortices. Here insular activation also presented a distinct activation for the LDX and PLA groups. Both these results (similar BOLD activations and insula) would come to corroborate the ones presented here. They would indicate that for this kind of smell ingestion of LDX was producing “normal” (understood as response from control group) odor function. A smell with a large emotional component.

In contrast to this, non-food related smells presented large changes with respect to volumes of activated voxels after LDX ingestion (Fig. 3). We could hypothesise that the involvement of more rational centers of the brain (mainly frontal structures) in drug-naive volunteers for non-food related odors, could imply that: First, they were more difficult to classify or interpretate; and second, they certainly did not elicit any kind of emotional response which we found associated to chocolate. This hypothesis is based on findings previously presented by this group (30). Still after ingestion of LDX Lavender and clove started presenting distinct activity in some detection centers of the olfactive center, these activations were mixed in the sense that some were stronger for the PLA group and other for the LDX. Therefore, these results are inconclusive. If figure 3 is observed, clove and lavender after LDx ingestion presented large increases of activations in pretty much the whole brain (frontal, temporal and occipital cortex as well as basal ganglia and olfactive centers). Differences as seen in Figure 2 (**B, C**) were only significant in temporal lobes for clove and occipital for lavender. Nevertheless LDX as an stimulant is having an effect of helping getting the whole brain involved in odor processing, when it was not happening for the PLA. This patter of activation was similar then to that of chocolate. So we con conclude by this pattern of behavior that LDX stimulates function in brain regions when smell was not eliciting any activity in drug-naïve people. To affirm that this activation was similar to those of non-ADHD patients a different and complete study would be necessary, but there is a trend of similarity for the chocolate flavor.

Authors do not think that any conclusion can be derived of the differences between activations due to food related smells (chocolate and clove) and non-food odors (lavender). This because of small sample size and lack of a more specific study.

Activations in smell detection regions of the brain were similar in magnitude across the PLA and LDX situations. Meanwhile, inhibitory regions of the brain such as the cingulate cortex and frontal lobe regions changed their activity pattern (different subregions in those areas were activated) and also showed changes in the intensity of activity (increasing with medication intake). As could be seen in Figure 1 and quantified in Table 1. The study of the effect of LDX on different flavors (Figures 2 & 3, Table 2), highlighted the fact that First, LDX increased brain activity in general when a flavor did not entice it on its own in the PLA group, and second, that this activity tended to be similar to that of healthy children even if a more thorough and conclusive work is needed to confirm this point. Because of all this, and as a conclusion to this section, it could be said that LDX first changed the inhibitory pathways of the brain, and second facilitated a complementary behavioral regulation mechanism run by the cerebellum, which regulated decision-making and impulsivity in motor and frontal structures.

## Author Contributions

Silvia S. Hidalgo Tobón: Obtained ethical permission for the study, performed measurements and discussed the results. Pilar Dies Suárez: Obtained ethical permission for the study and performed measurements. Eduardo Barragán Pérez: Analyzed data, discussed findings and helped write the manuscript. Javier M. Hernández López: Analyzed data, discussed findings and helped write the manuscript. Julio Flores: Analyzed data, discussed findings and helped write the manuscript. Benito de Celis Alonso: Analyzed data, discussed findings and helped write the manuscript. All authors fulfil the journal’s criteria of 1) substantial contributions to the work, or the acquisition, analysis, or interpretation of data for the work; 2) drafting the work or revising it critically; 3) final approval of the version to be published; and 4) agreement to be accountable for all aspects of the work.

## Data Availability of Data and Materials

All data and protocols used for this study will be available for researchers under petition to the corresponding author.

## Abbreviations

(LDX): Lisdexamfetamine,
(BED): binge eating disorder,
(ADHD): attention deficit hyperactivity disorder,
(fMRI): functional magnetic resonance imaging,
(PLA): placebo,
(PET): positron emission tomography,
(VMPFC): ventromedial prefrontal cortex,
(DLPC): dorsolateral prefrontal cortex recruitment,
(CGI test): Conners’ Global Index,
(BOLD): blood oxygenation level dependence,
(CSF): cerebrospinal fluid,
(MNI): Montreal Neurological Institute,
(FWE): family-wise error correction,
(ACC): anterior cingulate cortex,
(BA): Brodmann area,
(PCC): posterior cingulate cortex.

## Legends

**Table 1: Quantification of BOLD-fMRI results for all odor cues together.** Quantified data is shown from regions activated for the three contrasts presented in Figure 1. Data is presented for all olfactory stimuli together for the following contrasts: PLA, LDX and LDX– PLA. For each stimulus, first the number of activated voxels in each significant cluster is given. When a number of voxels is not followed in the following row by other number, it means that the same cluster was large enough to have different maximum points. After this are given the regions that these clusters were in. This is done showing both the corresponding AAL region and the BA in which they appear. In the last four columns, the statistical significance of the maximum voxel is presented, together with its MNI coordinates in mm. Brain regions that belong to the human olfaction system as described in introduction, are presented in italics for clarity.

**Table 2: Quantification of BOLD-fMRI results for each odor cue for the LDX – PLA contrast.** Quantified data is shown from regions activated for the three contrasts presented in Figure 2. Data is presented for all olfactory stimuli separately, for the following contrasts: chocolate LDX–PLA, clove LDX–PLA and lavender LDX–PLA.

## References

1. Kose S, Steinberg JL, Moeller FG, Gowin JL, Zuniga E, Kamdar ZN, et al. Neural correlates of impulsive aggressive behavior in subjects with a history of alcohol dependence. Behavioral neuroscience. 2015;129(2):183-96. Epub 2015/02/11.

2. Rusnakova S, Daniel P, Chladek J, Jurak P, Rektor I. The executive functions in frontal and temporal lobes: a flanker task intracerebral recording study. Journal of clinical neurophysiology: official publication of the American Electroencephalographic Society. 2011;28(1):30-5. Epub 2011/01/12.

3. Schag K, Schonleber J, Teufel M, Zipfel S, Giel KE. Food-related impulsivity in obesity and binge eating disorder--a systematic review. Obesity reviews: an official journal of the International Association for the Study of Obesity. 2013;14(6):477-95. Epub 2013/01/22.

4. Waxman SE. A systematic review of impulsivity in eating disorders. European eating disorders review: the journal of the Eating Disorders Association. 2009;17(6):408-25. Epub 2009/06/24.

5. Kalenscher T, Ohmann T, Gunturkun O. The neuroscience of impulsive and self-controlled decisions. International journal of psychophysiology: official journal of the International Organization of Psychophysiology. 2006;62(2):203-11. Epub 2006/07/11.

6. Evenden JL. Varieties of impulsivity. Psychopharmacology. 1999;146(4):348-61. Epub 1999/11/07.

7. McDonald V, Hauner KK, Chau A, Krueger F, Grafman J. Networks underlying trait impulsivity: Evidence from voxel-based lesion-symptom mapping. Human brain mapping. 2017;38(2):656-65. Epub 2016/09/27.

8. Whelan R, Conrod PJ, Poline JB, Lourdusamy A, Banaschewski T, Barker GJ, et al. Adolescent impulsivity phenotypes characterized by distinct brain networks. Nature neuroscience. 2012;15(6):920-5. Epub 2012/05/01.

9. Bear MF, Connors BW. Neuroscience exploring the brain: Lippincot williams & Wilkins; 2007.

10. Neville KR, Lewis B.H. The synaptic organization of the brain. Olfactory cortex. Oxford Univeristy Press 2004.

11. Subcommittee on Attention-Deficit/Hyperactivity D, Steering Committee on Quality I, Management, Wolraich M, Brown L, Brown RT, et al. ADHD: clinical practice guideline for the diagnosis, evaluation, and treatment of attention-deficit/hyperactivity disorder in children and adolescents. Pediatrics. 2011;128(5):1007-22. Epub 2011/10/18.

12. Biederman J, Krishnan S, Zhang Y, McGough JJ, Findling RL. Efficacy and tolerability of lisdexamfetamine dimesylate (NRP-104) in children with attention-deficit/hyperactivity disorder: a phase III, multicenter, randomized, double-blind, forced-dose, parallel-group study. Clinical therapeutics. 2007;29(3):450-63. Epub 2007/06/20.

13. Punja S, Shamseer L, Hartling L, Urichuk L, Vandermeer B, Nikles J, et al. Amphetamines for attention deficit hyperactivity disorder (ADHD) in children and adolescents. The Cochrane database of systematic reviews. 2016;2:CD009996. Epub 2016/02/05.

14. Barragán Pérez E, García Beristain JC, Hidalgo Gutiérrez R. Evaluation of the response of lisdexamfetamine in children and adolescents with ADHD. Salud Mental. 2018;41(6):6.

15. Coghill DR, Caballero B, Sorooshian S, Civil R. A systematic review of the safety of lisdexamfetamine dimesylate. CNS drugs. 2014;28(6):497-511. Epub 2014/05/03.

16. Fleck DE, Eliassen JC, Guerdjikova AI, Mori N, Williams S, Blom TJ, et al. Effect of lisdexamfetamine on emotional network brain dysfunction in binge eating disorder. Psychiatry research Neuroimaging. 2019;286:53-9. Epub 2019/03/25.

17. Shanmugan S, Loughead J, Nanga RP, Elliott M, Hariharan H, Appleby D, et al. Lisdexamfetamine Effects on Executive Activation and Neurochemistry in Menopausal Women with Executive Function Difficulties. Neuropsychopharmacology: official publication of the American College of Neuropsychopharmacology. 2017;42(2):437-45. Epub 2016/08/24.

18. Schulz KP, Krone B, Adler LA, Bedard AV, Duhoux S, Pedraza J, et al. Lisdexamfetamine Targets Amygdala Mechanisms That Bias Cognitive Control in Attention-Deficit/Hyperactivity Disorder. Biological psychiatry Cognitive neuroscience and neuroimaging. 2018;3(8):686-93. Epub 2018/04/18.

19. Biezonski D, Shah R, Krivko A, Cha J, Guilfoyle DN, Hrabe J, et al. Longitudinal magnetic resonance imaging reveals striatal hypertrophy in a rat model of long-term stimulant treatment. Translational psychiatry. 2016;6(9):e884. Epub 2016/09/07.

20. Saive AL, Royet JP, Plailly J. A review on the neural bases of episodic odor memory: from laboratory-based to autobiographical approaches. Frontiers in behavioral neuroscience. 2014;8:240. Epub 2014/07/30.

21. Wang J, Eslinger PJ, Smith MB, Yang QX. Functional magnetic resonance imaging study of human olfaction and normal aging. The journals of gerontology Series A, Biological sciences and medical sciences. 2005;60(4):510-4. Epub 2005/06/04.

22. Weismann M, Yousry I, Heuberger E, Nolte A, Ilmberger J, Kobal G, et al. Functional magnetic resonance imaging of human olfaction. Neuroimaging clinics of North America. 2001;11(2):237-50, viii. Epub 2001/08/08.

23. Kim YK, Hong SL, Yoon EJ, Kim SE, Kim JW. Central presentation of postviral olfactory loss evaluated by positron emission tomography scan: a pilot study. American journal of rhinology & allergy. 2012;26(3):204-8. Epub 2012/05/31.

24. Gogos JA, Osborne J, Nemes A, Mendelsohn M, Axel R. Genetic ablation and restoration of the olfactory topographic map. Cell. 2000;103(4):609-20. Epub 2000/12/07.

25. Al Ain S, Poupon D, Hetu S, Mercier N, Steffener J, Frasnelli J. Smell training improves olfactory function and alters brain structure. NeuroImage. 2019;189:45-54. Epub 2019/01/11.

26. Royet JP, Zald D, Versace R, Costes N, Lavenne F, Koenig O, et al. Emotional responses to pleasant and unpleasant olfactory, visual, and auditory stimuli: a positron emission tomography study. The Journal of neuroscience: the official journal of the Society for Neuroscience. 2000;20(20):7752-9. Epub 2000/10/12.

27. A.J.D. Crow, J.M. Janssen, K.L. Vickers, J. Parish-Morris, P.J. Moberg, D.R. Roalf. Olfactory Dysfunction in Neurodevelopmental Disorders: A Meta-analytic Review of Autism Spectrum Disorders, Attention Deficit/Hyperactivity Disorder and Obsessive-Compulsive Disorder. J Autism Dev Disord. 2020.

28. Ghanizadeh A, Bahrani M, Miri R, Sahraian A. Smell identification function in children with attention deficit hyperactivity disorder. Psychiatry investigation. 2012;9(2):150-3. Epub 2012/06/19.

29. Fuermaier ABM, Hupen P, De Vries SM, Muller M, Kok FM, Koerts J, et al. Perception in attention deficit hyperactivity disorder. Attention deficit and hyperactivity disorders. 2018;10(1):21-47. Epub 2017/04/13.

30. de Celis-Alonso B, Hidalgo-Tobon SS, Barragan-Perez E, Castro-Sierra E, Dies-Suarez P, Garcia J, et al. Different food odors control brain connectivity in impulsive children. CNS & neurological disorders drug targets. 2018. Epub 2018/11/06.

31. A.M. Herman, C. Hugo, Duka. T. Decreased olfactory discrimination is associated with impulsivity in healthy volunteers. Sci Rep. 2018;8(15584).

32. Godoy MD, Voegels RL, Pinna Fde R, Imamura R, Farfel JM. Olfaction in neurologic and neurodegenerative diseases: a literature review. International archives of otorhinolaryngology. 2015;19(2):176-9. Epub 2015/05/21.

33. El Fotoh W, El Naby SAA, Abd El Hady NMS. Autism Spectrum Disorders: The Association with Inherited Metabolic Disorders and Some Trace Elements. A Retrospective Study. CNS & neurological disorders drug targets. 2019;18(5):413-20. Epub 2019/06/19.

34. Pareek V, Nath B, Roy PK. Role of Neuroimaging Modality in the Assessment of Oxidative Stress in Brain: A Comprehensive Review. CNS & neurological disorders drug targets. 2019;18(5):372-81. Epub 2019/10/04.

35. Dileo JF, Brewer WJ, Hopwood M, Anderson V, Creamer M. Olfactory identification dysfunction, aggression and impulsivity in war veterans with post-traumatic stress disorder. Psychological medicine. 2008;38(4):523-31. Epub 2007/10/02.

36. Rolls ET. Taste, olfactory and food texture reward processing in the brain and obesity. International journal of obesity. 2011;35(4):550-61. Epub 2010/08/04.

37. Best M, Williams JM, Coccaro EF. Evidence for a dysfunctional prefrontal circuit in patients with an impulsive aggressive disorder. Proceedings of the National Academy of Sciences of the United States of America. 2002;99(12):8448-53. Epub 2002/05/30.

38. Villafuerte G, Miguel-Puga A, Arias-Carrión O. Continuous Theta Burst Stimulation Over the Right Orbitofrontal Cortex Impairs Conscious Olfactory Perception. Front Neurosci,. 2019. Epub 05/06/2019.

39. Yeomans MR. Olfactory influences on appetite and satiety in humans. Physiology & behavior. 2006;89(1):10-4. Epub 2006/05/23.

40. Alonso Bde C, Hidalgo Tobon S, Dies Suarez P, Garcia Flores J, de Celis Carrillo B, Barragan Perez E. A multi-methodological MR resting state network analysis to assess the changes in brain physiology of children with ADHD. PloS one. 2014;9(6):e99119. Epub 2014/06/20.

41. Guerrero Arenas C, Hidalgo Tobon SS, Dies Suarez P, Barragan Perez E, Castro Sierra E, Garcia J, et al. Strategies for tonal and atonal musical interpretation in blind and normally sighted children: an fMRI study. Brain and behavior. 2016;6(4):e00450. Epub 2016/04/12.

42. Hayasaka S, Peiffer AM, Hugenschmidt CE, Laurienti PJ. Power and sample size calculation for neuroimaging studies by non-central random field theory. NeuroImage. 2007;37(3):721-30. Epub 2007/07/31.

43. Yechiam E, Goodnight J, Bates JE, Busemeyer JR, Dodge KA, Pettit GS, et al. A formal cognitive model of the go/no-go discrimination task: evaluation and implications. Psychological assessment. 2006;18(3):239-49. Epub 2006/09/07.

44. Jonkman LM, Lansbergen M, Stauder JE. Developmental differences in behavioral and event-related brain responses associated with response preparation and inhibition in a go/nogo task. Psychophysiology. 2003;40(5):752-61. Epub 2003/12/31.

45. Slobodin O, Cassuto H, Berger I. Age-Related Changes in Distractibility: Developmental Trajectory of Sustained Attention in ADHD. Journal of attention disorders. 2018;22(14):1333-43. Epub 2015/03/21.

46. Conners CK, Sitarenios G, Parker JD, Epstein JN. The revised Conners’ Parent Rating Scale (CPRS-R): factor structure, reliability, and criterion validity. Journal of abnormal child psychology. 1998;26(4):257-68. Epub 1998/08/13.

47. Hellman TM, Small FH. Characterization of the odor properties of 101 petrochemicals using sensory methods. J Air Pollut Control Assoc. 1974;24(10):3.

48. Dravnieks AT, Masurat S, Lamm RA. Hedonics of odors and odor descriptors. Air Poll Cont Ass. 1984;37(4).

49. Whitfield-Gabrieli S, Nieto-Castanon A. Conn: a functional connectivity toolbox for correlated and anticorrelated brain networks. Brain connectivity. 2012;2(3):125-41. Epub 2012/05/31.

50. Yan CG, Wang XD, Zuo XN, Zang YF. DPABI: Data Processing & Analysis for (Resting-State) Brain Imaging. Neuroinformatics. 2016;14(3):339-51. Epub 2016/04/15.

51. Yousem DM, Maldjian JA, Siddiqi F, Hummel T, Alsop DC, Geckle RJ, et al. Gender effects on odor-stimulated functional magnetic resonance imaging. Brain research. 1999;818(2):480-7. Epub 1999/03/20.

52. de Celis Alonso B, Sergeyeva M, Brune K, Hess A. Lateralization of responses to vibrissal stimulation: connectivity and information integration in the rat sensory-motor cortex assessed with fMRI. NeuroImage. 2012;62(3):2101-9. Epub 2012/06/06.

53. Zucco G, Tressaldi PE. Hemispheric differences in odor recognition. Cortex. 1988;25:8.

54. Henkin RI, L.M. L. Lateralization of Brain Activation to Imagination and Smell of Odors Using Functional Magnetic Resonance Imaging (fMRI): Left Hemispheric Localization of Pleasant and Right Hemispheric Localization of Unpleasant Odors Journal of Computer Assisted Tomography 2001;25(4):21.

55. Mohamed SM, Borger NA, Geuze RH, van der Meere JJ. Brain lateralization and self-reported symptoms of ADHD in a population sample of adults: a dimensional approach. Frontiers in psychology. 2015;6:1418. Epub 2015/10/07.

56. Garcia-Cabezas MA, Barbas H. A direct anterior cingulate pathway to the primate primary olfactory cortex may control attention to olfaction. Brain structure & function. 2014;219(5):1735-54. Epub 2013/06/26.

57. Ding WN, Sun JH, Sun YW, Chen X, Zhou Y, Zhuang ZG, et al. Trait impulsivity and impaired prefrontal impulse inhibition function in adolescents with internet gaming addiction revealed by a Go/No-Go fMRI study. Behavioral and brain functions: BBF. 2014;10:20. Epub 2014/06/03.

58. Bush G, Frazier JA, Rauch SL, Seidman LJ, Whalen PJ, Jenike MA, et al. Anterior cingulate cortex dysfunction in attention-deficit/hyperactivity disorder revealed by fMRI and the Counting Stroop. Biological psychiatry. 1999;45(12):1542-52. Epub 1999/06/22.

59. Leech R, Sharp DJ. The role of the posterior cingulate cortex in cognition and disease. Brain: a journal of neurology. 2014;137(Pt 1):12-32. Epub 2013/07/23.

60. Liddle EB, Hollis C, Batty MJ, Groom MJ, Totman JJ, Liotti M, et al. Task-related default mode network modulation and inhibitory control in ADHD: effects of motivation and methylphenidate. Journal of child psychology and psychiatry, and allied disciplines. 2011;52(7):761-71. Epub 2010/11/16.

61. Castellanos FX, Margulies DS, Kelly C, Uddin LQ, Ghaffari M, Kirsch A, et al. Cingulate-precuneus interactions: a new locus of dysfunction in adult attention-deficit/hyperactivity disorder. Biological psychiatry. 2008;63(3):332-7. Epub 2007/09/25.

62. Peterson BS, Potenza MN, Wang Z, Zhu H, Martin A, Marsh R, et al. An FMRI study of the effects of psychostimulants on default-mode processing during Stroop task performance in youths with ADHD. The American journal of psychiatry. 2009;166(11):1286-94. Epub 2009/09/17.

63. Moers-Hornikx VM, Sesia T, Basar K, Lim LW, Hoogland G, Steinbusch HW, et al. Cerebellar nuclei are involved in impulsive behaviour. Behavioural brain research. 2009;203(2):256-63. Epub 2009/05/20.

64. Blackwood N, Ffytche D, Simmons A, Bentall R, Murray R, Howard R. The cerebellum and decision making under uncertainty. Brain research Cognitive brain research. 2004;20(1):46-53. Epub 2004/05/08.

65. Schmahmann JD. The cerebellum and cognition. Neuroscience letters. 2019;688:62-75. Epub 2018/07/13.

66. Small DM, Jones Gotman M, Zatorre RJ, Petrides M, Evans AC. Flavor processing: More than the sum of its parts. Neuroreport 1997;22(8):4.

67. Savic I. Processing of odorous signals in humans. Brain research bulletin. 2001;54(3):307-12. Epub 2001/04/05.

68. Forstmann BU, van den Wildenberg WP, Ridderinkhof KR. Neural mechanisms, temporal dynamics, and individual differences in interference control. Journal of cognitive neuroscience. 2008;20(10):1854-65. Epub 2008/03/29.

